# Genome assembly of the JD17 soybean provides a new reference genome for Comparative genomics

**DOI:** 10.1101/2021.11.23.469778

**Authors:** Xinxin Yi, Jing Liu, Shengcai Chen, Hao Wu, Min Liu, Qing Xu, Lingshan Lei, Seunghee Lee, Bao Zhang, Dave Kudrna, Wei Fan, Rod A. Wing, Chunyan Yang, Mengchen Zhang, Jianwei Zhang, Xuelu Wang, Nansheng Chen

## Abstract

Cultivated soybean (*Glycine max*) is an important source for protein and oil. Many elite cultivars with different traits have been developed for different conditions. Each soybean strain has its own genetic diversity, and the availability of more high-quality soybean genomes can enhance comparative genomic analysis for identifying genetic underpinnings for its unique traits. In this study, we constructed a high-quality *de novo* assembly of an elite soybean cultivar Jidou 17 (JD17) with chromsome contiguity and high accuracy. We annotated 52,840 gene models and reconstructed 74,054 high-quality full-length transcripts. We performed a genome-wide comparative analysis based on the reference genome of JD17 with three published soybeans (WM82, ZH13 and W05), which identified five large inversions and two large translocations specific to JD17, 20,984 - 46,912 PAVs spanning 13.1 - 46.9 Mb in size, and 5 - 53 large PAV clusters larger than 500kb. 1,695,741 - 3,664,629 SNPs and 446,689 - 800,489 Indels were identified and annotated between JD17 and them. Symbiotic nitrogen fixation (SNF) genes were identified and the effects from these variants were further evaluated. It was found that the coding sequences of 9 nitrogen fixation-related genes were greatly affected. The high-quality genome assembly of JD17 can serve as a valuable reference for soybean functional genomics research.

## Introduction

Soybean (*Glycine max*) is an important crop for protein and dietary oil and is ranked the fourth largest crop in production in the world. The current soybean reference genome, which was based on the Williams 82 (WM82) line (Schmutz et al. 2010a) has greatly enhanced the identification of genes underlying important traits and facilitated research on the function and expression of soybean genes.

Recent studies using high-throughput sequencing has revealed extensive genetic diversities in soybean (Zhou et al. 2015). Pan-genome study on wild and cultivated soybeans has uncovered numerous genetic differences among soybean strains (Liu et al. 2020), suggesting that a single reference genome is inadequate for representing the genetic richness of soybean lines.

The latest progress in sequencing technologies has greatly advanced the ability to construct high-quality genome assemblies with chromosome-level continuity with dramatically reduced cost (Risse et al. 2015; Deschamps et al. 2018). The application of Iso-Seq protocol has enhanced genome annotation (Jiao et al. 2017; Li et al. 2017; Jiang et al. 2017; Magrini et al. 2018). Benefiting from improved technologies, more and more reference genomes of various soybean materials have been sequenced and reported, including the Chinese cultivar ZH13(Shen et al. 2019), three reference genomes (*Glycine max* WM82v4, *Glycine max* Lee and *Glycine soja* PI 483463)(Valliyodan et al. 2019), the wild strain W05 genome(Xie et al. 2019), the pan-genome constructed from 26 different soybean species, and the recently published Korean Hwangkeum genome and eight soybean genomes(Chu et al. 2021).

JD17 is a major soybean cultivar in the Huang-Huai-Hai region of China, and it is also the main soybean variety recognized by the Ministry of Agriculture since 2010. It is the offspring of Hobbit (maternal parent) and Zao 5241 [7476 × 7527-1-1 (Yanli×Williams)] (paternal parent) (Qin et al. 2014), and is famous for its lodging resistance, high yield, and strong adaptability (Zhao et al. 2013; Zhao et al. 2015). The goal of this project was to construct a high-quality genome assembly for JD17, and apply the whole genome sequence to identify genetic basis of important traits at the molecular level. In 2019, nodulation and symbiotic nitrogen fixation in legume were summarized and aggregated(Roy et al. 2020), but we still have many unknowns about the reasons for phenotypic differences in rhizobia number, morphology, and nitrogen fixation capacity in different strains of soybean.

In this study, we extracted genome DNA from developing underground tissues of JD17, and used PacBio single-molecule real-time (SMRT) sequencing and Hi-C mapping technologies to construct a high-quality soybean reference genome. After being annotated more completely, the JD17 pseudomolecules were used to identify structural differences in a comparative analysis with three published soybean reference genomes. We identified JD17-specific PAVs and a large number of SNPs and Indels, and also resolved the influence of these variants on the coding structure of nitrogen fixation-related genes.

## Materials and methods

### Plant and Sample preparation

Soybean seeds of *Glycine max* cv. JD17 used in this study were from Hebei Academy of Agricultural and Forestry Sciences. The seeds are planted and extracted in Xuelu Wang’s Laboratory in Huazhong Agricultural University. The seeds were sterilized with chlorine gas (5 ml of 32% (w/w) HCl to 100 ml 4–5% (wt/vol) sodium hypochlorite in a beaker) for 15 h (Kereszt et al. 2007) and then left in a sterile hood for 2 h. The sterilized seeds were sown in growth bottles filled with sand after being soaked in sterile Milli-Q water for 30 mins and watered with sterile Fahraeus solution (FÅHRAEUS 1957) containing 2 mM KNO_3_. Seeds were grown in a growth chamber (light) at 28℃ and 8 h (dark) at 23℃ with 60% humidity for 16 h.

### RNA preparation and Sequencing

Underground tissues of inoculation and uninoculation from the 9 timepoints (1, 4, 6, 8, 10, 15, 20, 25 and 30 day post inoculation, dpi) were collected and used for RNA extraction respectively. After that, nine RNA samples of inoculation and uninoculation were mixed equally as a sample, respectively. In addition, we also selected different tissues including root, nodule, stem, leaf, pod, seed and flower for mixed RNA-Seq. All the RNA was extracted by TRIzol reagent (Invitrogen 15596026). we performed RNA-Seq on illumina platform and produced approximately 10 Gb raw data with 150 bp pair-end reads.

### Whole-genome sequencing using SMRT technology

10-dpi root tissue of plants was used for SMRT whole-genome sequencing. Underground tissues were collected for genomic DNA preparation with modified CTAB method (Bergman and Quesneville 2007). Using 97 ug DNA, PacBio sequencing libraries were produced following manufacturers protocols as described for the Greater than 30kb-SMRTbell Libraries Needle Shearing (SMRTbell Template Prep Kit 1.0) with Blue Pippin size selections (Sage Science, http://www.sagescience.com/), and the SMRTbell libraries were constructed through Pacific Biosciences SMRTbell Template Prep Kit 1.0 (http://www.pacb.com/). SMRT sequencing was performed on a PacBio RSII instrument using P6/C4 sequencing chemistry (DNA/Polymerase Binding Kit P6) and 6hr movies. We used a total of 118 SMRT cells and produced 127.3 Gb of raw data with an average subread length of 15 kb (Table S1). At the same time, we also used a part of DNA samples for resequencing on the Illumina HiSeq 2500 platform for 150 bp paired end reads, with a sequencing depth of approximately 45X, for a total of 44.5Gb. These data are mainly used for the evaluation of genome size and its heterozygosity, post-assembly error correction and genome quality assessment.

### Hi-C library construction and sequencing

For samples used for Hi-C assisted assembly, leaves fixed in 1% (volume / volume) formaldehyde were used for library construction. Cell lysis, chromatin digestion, proximity ligation treatment, DNA recovery and subsequent DNA manipulation were performed as previously described (Lieberman-Aiden et al. 2009). The restriction enzyme used in chromatin digestion is Mbol. Finally, the Hi-C library was sequenced on the Illumina HiSeq X 10 platform for 150 bp paired end reads, with a sequencing depth of approximately 150X, for a total of 152.0 Gb.

### *De novo* genome assembly of JD17

To perform *de novo* assembly of the JD17, we combined three different assemblers, including CANU (Koren et al. 2017) (v1.4), FALCON (https://github.com/PacificBiosciences/FALCON-integrate) and HGAP4 (SMRT Link v 5.0.1.9585) (Table S2). The main assembly was performed on whole SMRT sequenced long reads. All assembly softwares were performed with a presumed 1-Gb genome size. If not specified, all programs in our study were run with default parameters. CANU was run with ‘errorRate = 0.013’, and FALCON was run with ‘length_cutoff = -1’ for initial mapping of seed reads for the error-correction phase (Figure S1). For a better FALCON assembly, we additionally optimized parameters as ‘DBsplit = -x500, --s400, pa_HPCdaligner = -v -B128 -t16 -e.70 -l1000 -s1000 -T8 -M24, ovlp_HPCdaligner = -v -B128 -t32 -h60 -e.96 -l500 -s1000 -T8 -M24 and overlap_filtering = --max_diff 60 --max_cov 60 --min_cov 2’. The stats of three initial assemblies were show in Table S2. We subsequently used the CANU assembly as the working set because it was better able to phase the diploid genome as well as indicating better contiguity. Subsequently, the draft assembly was polished twice using Quiver (SMRT Link v 5.0.1.9585) and finally corrected using Illumina short reads with Pilon (Figure S2). Then non-soybean contigs were filtered by aligning them to NT library using blastn.

### Continuation and connection of contigs

Due to the complementarity among three different assembly results from CANU (Koren et al. 2017), FALCON and HGAP4, we optimized our assembly results using the GPM (Zhang et al. 2016) pipeline to extend and connect contigs for better contiguity. Firstly, GPM loaded WM82 as a reference genome, and CANU assembly as the back-bone contigs. These contigs were ordered and located on chromosome based on WM82 (Glycine_max_v2.0) using blastn. This step produced a draft chromosome assembly, as JD17 v0.1. Secondly, we loaded FALCON assembly result and aligned with CANU assembly using blastn (Camacho et al. 2009a), and got the potential overlapping relationships between CANU and FALCON contigs. Then CANU contigs were extended or connected by FALCON contigs as needed to produce the JD17 v0.2 assembly. Thirdly, we repeated the above steps with HGAP4 assembly and the JD17 v0.3 was generated. Fourthly, we loaded contigs from an assembly performed with the longest 70 Gb PacBio reads (extracted from the total 120 Gb sorted raw reads) using CANU and repeated the above editing steps to reach the JD17 v0.4. Finally, the contigs of chloroplast and mitochondrial sequences were identified and removed from JD17 v0.4 (Table S3). At editing steps, the JD17 v0.4 was re-polished using Quiver over twice iterations and corrected using Illumina short reads with Pilon (Walker et al. 2014). The finalized JD17 genome assembly (named as Glycine_max_JD17v1.0) was 995.0 Mb in size, with a contig N50 of 18.0 Mb (Table 1). The assembly processing details are shown in the Figure S3.

**Table 1.** Assembly statistics of Glycine_max_JD17 (JD17), Glycine_max_v4.0 (WM82), Gmax_ZH13 (ZH13) and W05.

| Assembly Feature | JD17 | WM82 | ZH13 | W05 |
| --- | --- | --- | --- | --- |
| Estimated genome size (by K-mer analysis) (Mb) | 1,109 | 1,115 | - | - |
| Number of contigs | 446 | 9,200 | 1,528 | 1,870 |
| Total size of contigs (Mb) | 995.0 | 952.5 | 1,007 | 998.6 |
| Longest contig (Mb) | 31.8 | - | - | - |
| Number of contigs > 1 Mb | 97 | - | - | - |
| Number of contigs > 10 Mb | 39 | - | - | - |
| N50 contig length (Mb) | 18.0(PacBio) | 0.4 | CANU:2.9<br>+Bionno:18.0<br>+Hi-C:22.6 | 3.3 |
| L50 contig count | 21 | 649 | 66 | 58 |
| **Anchored contigs** |  |  |  |  |
| Number of chromosomes | 20 | 20 | 20 | 20 |
| Number of contigs | 411 | - | - | 772 |
| Total size (Mb) | 965.8 | 978.4 | 1,011.2 | 1,013.2 |
| Number of gaps | 391 | 7,221 | 448 | 750 |

The assessment of genomic heterozygosity and size is using the Genomic Character Estimator program (gce v 1.0.0, ftp://ftp.genomics.org.cn/pub/gce), and the heterozygous ratio based on kmer individuals is 0.029, and the corrected estimation of genome size is about 1.11 Gb.

To anchor hybrid contigs into chromosome, the Hi-C sequencing data were aligned into contigs using bwa. According to the orders and orientations provided by the alignment, those contigs were clustered into chromosomes by ALLHiC v0.9.8 (Zhang et al. 2019) with default params. According to the ALLHiC groups and assembly results create hic files, manual correction and validation were also performed by drawing contact maps with juicerbox (Durand et al. 2016). The genome assembly was finalized after this correction step (Table S3).

### Quality assessment of JD17 genome assembly

To assess the quality of the Glycine_max_JD17v1.0 assembly, we used our 65 Gb resequencing data. First, by aligning all reads to the assembly with BWA-MEM in BWA (Li 2014), the mapping rate is over 98.8% and the coverage was over 99.65%, which shows the consistency between the assembly and reads. By using the GATK tools (McKenna et al. 2010; DePristo et al. 2011) for SNPs calling with JD17 resequencing data, we found 78,033 SNP, of which only 8,000 were homozygous, indicating that the JD17 genome has an accuracy of over 99.999%. The completeness of the assembly was estimated by BUSCO with default parameters.

### Annotation of TE and ncRNA sequences

To investigate the JD17 genome sequence features, we identified transposable elements (TEs) and other repetitive elements by RepeatMasker (Bergman and Quesneville 2007). MITEs (miniature inverted transposable elements) were collected by MITE-Hunter (Han and Wessler 2010) with all default parameters. In order to get as much reliable LTR (long terminal repeat) retrotransposons information as possible, we used the LTR_retriever (Ou and Jiang 2018) analysis process, which integrates the output of LTR_FINDER (Xu and Wang 2007) and LTRharverst tools in GenomeTools (Gremme et al. 2013). Masking sequence with RepeatMasker (version 4.0.8) (http://www.repeatmasker.org/) based on MITEs and LTR library that has been identified. The other tandem repeats were identified by constructing a *de novo* repeat library using Repeatmodeler (version 1.0.11) (http://www.repeatmasker.org/). RepeatMasker was run against the genome assembly again, with all above library as the query library.

Non-coding RNAs were predicted by the Infernal program using default parameters (Nawrocki and Eddy 2013) and comparing the similarity of secondary structure between the JD17 genome sequence and Rfam (Nawrocki et al. 2014) (v12.0) database.

### Annotation of protein-coding genes

We performed gene calling analysis with Exonerate (Slater and Birney 2005), Trinity (Grabherr et al. 2011) and PASA (Haas et al. 2003), by using multi-sourced EST and protein sequences as evidences (including nonredundant soybean EST / Iso-seq (<u>SRX7016448</u>) (Chu et al. 2021) / protein sequences from NCBI, assembled JD17 RNA-Seq from mixed samples from underground samples, WM82 transcripts and proteins, *Lotus japonicus* and *Medicago truncatula* protein sequences), AUGUSTUS (Hoff and Stanke 2013) and GENEMARK (Bruna et al. 2020) as *ab initial* gene predictors, and a customized repeat library for RepeatMasker (Bergman and Quesneville 2007; Saha et al. 2008).

The RNA-Seq data was *de novo* assembled using Trinity to obtain the assembled cDNA sequence, and annotated with the PASA tool along with the non-redundant isoforms sequence from Iso-Seq. The annotation results will be used for AUGUSTUS model training and prediction. The WM82 protein sequence will be used for GENEMARK model training and prediction. Protein sequences of *Arabidopsis thaliana, Medicago sativa, Lotus corniculatus* and three different strains of soybean (WM82, ZH13, W05) were used in the Exonerate de-prediction protein-coding gene models. In order to obtain more accurate and complete annotation results, EVM was called to integrate the gene model prediction results from AUGUSTUS, GENEMARK, Exonerate and PASA. After all, the PASA software is used to update these annotation results.

Gene functions were inferred according to the best match of the alignments to the National Center for Biotechnology Information (NCBI) Swiss-Prot(Boeckmann et al. 2003) protein databases using BLASTP (ncbi blast v2.6.0+)(Altschul et al. 1997; Camacho et al. 2009b) and the Kyoto Encyclopedia of Genes and Genomes (KEGG) database(Kanehisa et al. 2012) with an E-value threshold of 1E-5. Gene Ontology (GO)(Ashburner et al. 2000) IDs for each gene were obtained from Blast2GO(Conesa and Gotz 2008).

### Genome-wide rearrangement and SV detection

To identify large-scale synteny among the four soybean lines, we created a genome-wide alignment using the Mauve aligner with the progressiveMauve algorithm (Darling et al. 2010) with default parameters: default seed weights, determination of LCBs (minimum weights = default), and full alignment with iterative refinement (Chakraborty et al. 2021).

The PAV sequences, PAV clusters and PAV genes between JD17 and the three genomes (WM82, ZH13, and W05) were identified using the sliding window method described in the 2018 publication by Sun et al (Sun et al. 2018), with the same slide window size, alignment software and parameters as theirs.

We aligned WM82, ZH13 and W05 to JD17 using MUMmer 4.0 (NUCmer-maxmatch) (Marcais et al. 2018). MUMmer alignments were processed using SyRI (Synteny and Rearrangement Identifier) (Goel et al. 2019) (https://github.com/schneebergerlab/syri commit 3f16e01), which identifies syntenic regions, chromosome rearrangement events such as inverson and translocation, and also identified SNPs, Indel, Copygains and Copylosses between chromosomes. The annotations of JD17 were indexed using the SnpEff tool for construction, and then SNPs and Indels identified genome-wide were annotated to finally find genes subject to large genetic variation.

### Identification and structural variation analysis of SNF genes

Identification of SNF gene sequences based on the literature published by Roy et al. in 2020 (Roy et al. 2020). These sequences were first aligned to the protein sequences of JD17, WM82, ZH13 and W05 by blastp tool and filtered by coverage and identity greater than 40% to obtain possible nitrogen fixation related genes. These filtered genes were then subjected to clustering analysis by the OrthoFinder tool (Emms and Kelly 2019). The SNF genes in the four soybeans were finally identified by determining the gene cluster where the published SNF genes were located. To obtain the expansion and contraction of SNF genes, these single-copy SNF genes were further used to construct an evolutionary tree by RAxML tool (Stamatakis 2014) and finally combined with café (De Bie et al. 2006) for the analysis of expansion and contraction.

Then the SNPs and Indels loci between JD17 and them were combined to assess the protein encoding affected SNF genes. These genes (and gene sequences from the article) were then aligned using mafft(Nakamura et al. 2018) software, constructing an evolutionary tree based on fasttree(Price et al. 2010) tools, and manually checking the evolutionary tree manually to determine that these genes whose structure was affected were homologous (protein-coding genes on the same chromosome; on the same branch of the evolutionary tree, and then expanded if genes of the species were not present on that branch).

## Results and discussion

### High quality genome assembly and annotation

Genome assembly using PacBio SMRT data and Hi-C data (detail in methods) resulted in a JD17 genome assembly with 411 contigs anchored to 20 pseudomolecules, with a total size of 965.8Mb and contigs N50 up to 18.0 Mb (Table 1), accounting for 97.1% of the total contigs (Figure 1). The genome accuracy rate was evaluated as 99.999% by using JD17 resequencing data and GATK tools (McKenna et al. 2010; DePristo et al. 2011) (See methods). BUSCO score of the assembly was 93.2% (Simão et al. 2015) (Table S4), indicating that the completeness of JD17 was higher than that of the WM82, ZH13, and W05 genomes (Shen et al. 2018; Xie et al. 2019).

**Figure 1.**
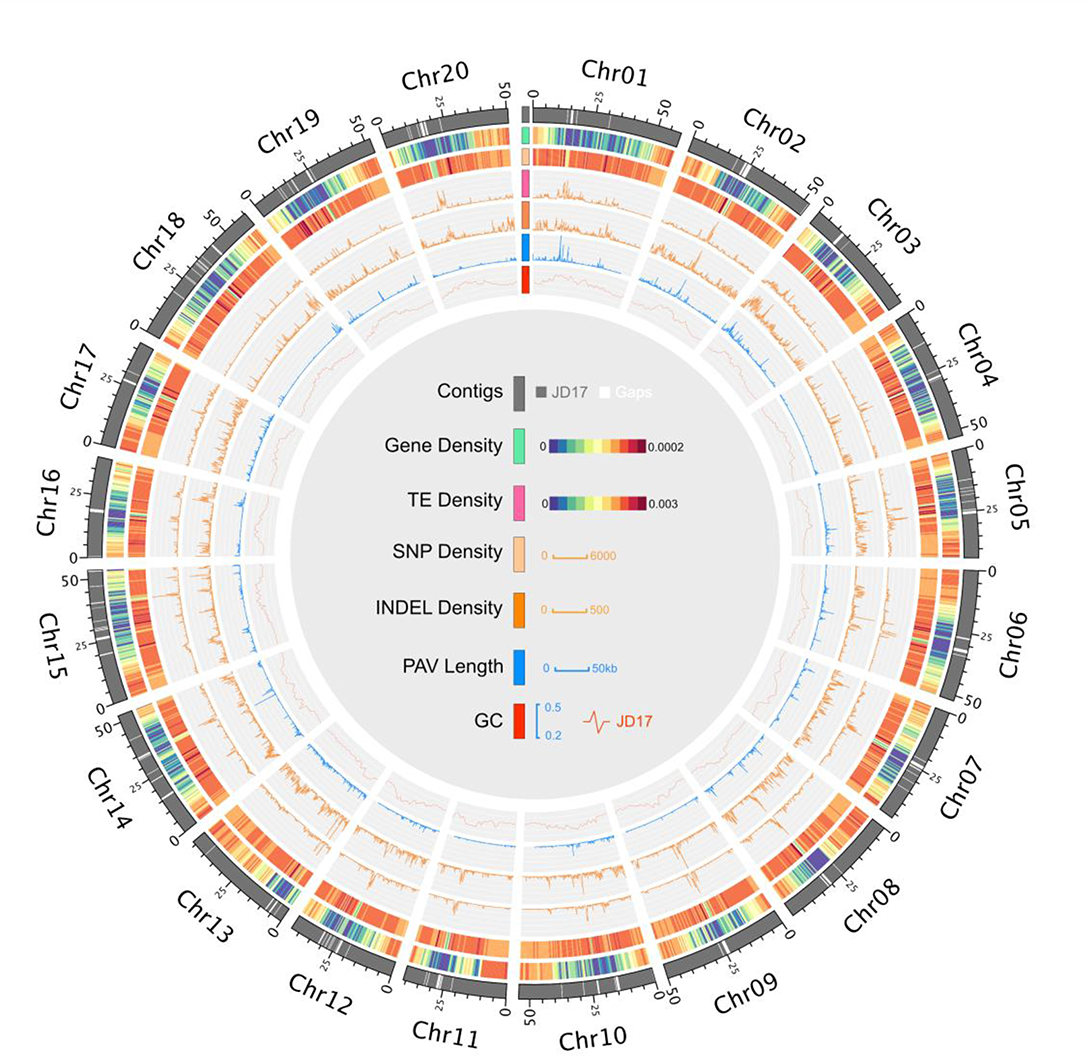
Overview of the JD17 reference genome. Tracks from outer to inner circles indicate: the chromosome of the genome; the gene density map; the repeat sequence density map; density distribution of SNPs between JD17 and WM82 ; density distribution of InDel between JD17 and WM82. PAV length distribution between JD17 and WM82; GC content of JD17.

In addition to the nuclear genome, we also constructed the full-length genomes of the soybean chloroplast (152.2 Kb) and the complete genome of *Bradyrhizobium japonicum* USDA 110 (9.1 Mb), which are consistent with previously reported results (Figure S4)(Saski et al. 2005; Kaneko et al. 2002).

We identified 52,840 protein-coding genes and 74,054 full-length transcripts in the JD17 genome (Table 2). The average length of mRNA, 5’UTR, CDS and 3’UTR were 4,465 bp, 302 bp, 1,183 bp and 487 bp, respectively. Our gene annotation is relatively in agreement with the other three genomes. After searching with existing databases and conserved structural domains for functional annotation, a total of 72,529 (97.94%) transcripts have known domains or functions, which suggested that high-confidence annotation of JD17 genome was perfomed.

**Table 2.**
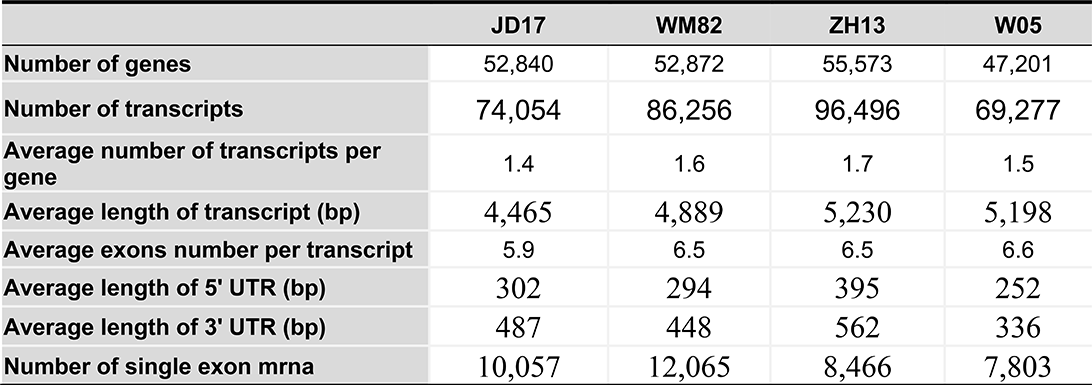
Comparison of genome annotation of JD17, WM82, ZH13 and W05.

|  | **JD17** | **WM82** | **ZH13** | **W05** |
| --- | --- | --- | --- | --- |
| **Number of genes** | 52,840 | 52,872 | 55,573 | 47,201 |
| **Number of transcripts** | 74,054 | 86,256 | 96,496 | 69,277 |
| **Average number of transcripts per gene** | 1.4 | 1.6 | 1.7 | 1.5 |
| **Average length of transcript (bp)** | 4,465 | 4,889 | 5,230 | 5,198 |
| **Average exons number per transcript** | 5.9 | 6.5 | 6.5 | 6.6 |
| **Average length of 5' UTR (bp)** | 302 | 294 | 395 | 252 |
| **Average length of 3' UTR (bp)** | 487 | 448 | 562 | 336 |
| **Number of single exon mrna** | 10,057 | 12,065 | 8,466 | 7,803 |

### Transposable elements

We identified 580.2Mb (58.30% of the JD17 genome) repeat elements in JD17 genome (Figure 2), which contain 1,138,787 intact transposable elements (TEs), including 734,061 class I RNA retrotransposons and 293,279 class II DNA transposons (Table S5).

**Figure 2.**
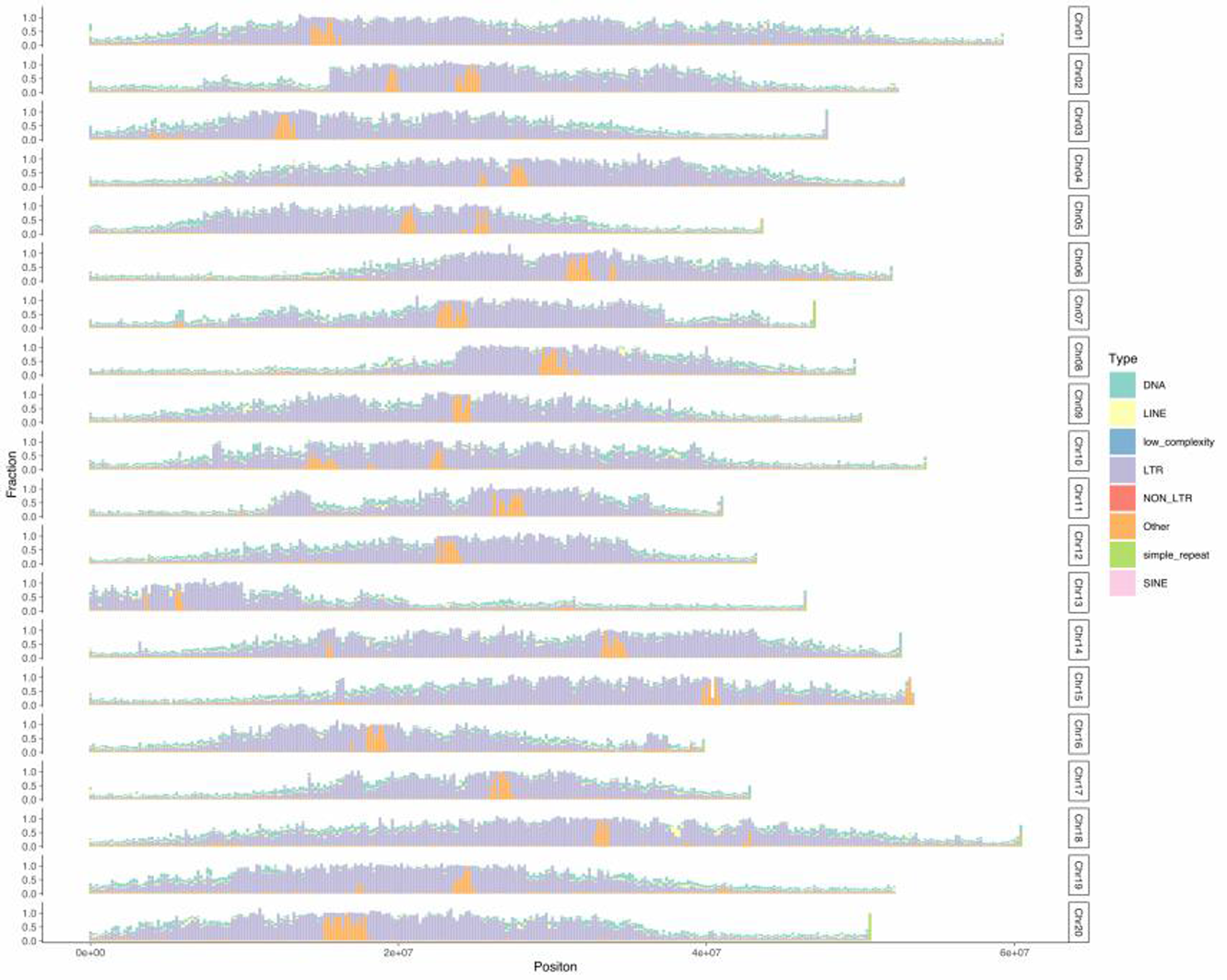
Chromosome distribution map and percentage of different types of TEs. Chromosomes were split into 200kb bins without overlap, and the percentage of major types of TE elements (include DNA: DNA transposons, LINEs: long interspersed nuclear elements, Low complexity, LTR: long terminal repeat retrotransposons, NON LTR: Non-long terminal repeat retrotransposons, other: simple repeat, SINE: Short interspersed nuclear elements) in each bin was counted.

Among the genomic sequences of JD17, the highest propotion of repetitive sequences is the long terminal repeat (LTR) retrotransposon, and a total of 631,056 LTR elements were identified, totaling 397.58 Mb, which is about 39.95% of the whole genome. Further by assessing the distribution and insertion time of LTRs, we found that the LTR enrichment was all in close proximity to the centromeric region, fewer in the gene enrichment region, and that the amplification of LTR retrotransposons in soybean occurred mainly within the past 1.5 million years and were all relatively young (Figure S5). When comparing the activity of LTR elements (Figure S6), Copia elements appear to be increasingly active and persistent over the past 3 million years. Similarly, Gypsy elements were relatively active during this period. In fact, between 1.4 and 3 million years, their activity levels do not differ much, but since 1.4 million years, the Gypsy element has become less active compared to the Copia element, while the unknown element has also become active over the last 3 million years, but at a much lower level compared to both the Gypsy and Copia elements. Based on the timing of the recent occurrence of WGD in soybean (∼13 Mya)(Schmutz et al. 2010b), it appears that these LTR insertions and expansions are different from gene duplications, but occur mainly after WGD.

To compare the differences in the composition and distribution of repetitive sequences between them, we annotated the repetitive sequences of other three genomes using the same approach. The results indicates that the composition of their TEs is essentially the same, except that WM82 has slightly less LTR content. (Figure 2, Table S5 and Figure S7-9). Meanwhile, we found some blank regions in the genomes of W82 and W05 in the distribution of TEs, both of which are N sequences in the genome sequence. To determine whether these blank regions are the centromeric regions, we identified the centromeric regions of the JD17 genome by using the soybean-specific centromeric satellite repeats sequences (CentGm-1 and CentGm- 2)(Gill et al. 2009). The evidence suggests that the location of JD17 centromeric regions is consistent with that of the blank regions in WM82 and W05 (Figure 2, Figure S10). In general, the assemblies of JD17 and ZH13 are more complete in the centromeric regions, and the TEs of W05 has slightly richer components than the other strains.

### Genomic rearrangements

To explore how many and how large scale discordant regions of genetic mapping exist between the JD17 reference genome and the three published genomes, we performed a genome-wide comparative analysis. When the pseudochromosomes of JD17 were aligned to the other pseudochromosomes of WM82, ZH13 and W05, a total of 3,727 syntenic blocks were identified (Figure S11), of which 1,368 were common syntenic blocks among the four strains (Table S6). Approximately 88.51% of the JD17 genome sequence shares the syntenic blocks with 88.12% of WM82, 86.10% of ZH13, and 81.88% of W05. We found a total of 251 syntenic blocks that exist between Wm82, ZH13 and W05, but not in JD17, with a size between 3.87-5.27Mb. Similarly, there are 113 such syntenic blocks between JD17, WM82 and ZH13, but not in W05, and the size is between 2.10-3.33Mb. The size of the specific sequence in each of the four strains (that is, the sequence that does not match any other strains) is between 82.58-119.37Mb, of which the wild-type W05 has the most specific sequences and the W82 has the least. This may be due to the loss of sequences in cultivars during the long domestication process of artificial selection.

The chromosomal differences between JD17 and the other three strains were further observed by comparative genome analysis. We observe that compared to WM82 and ZH13, JD17 has slightly more Copygains, 4.0Mb and 3.8Mb respectively, but W05 has more Copygains compared to JD17, with a combined total of about 7.1Mb (Table S7). We also found 56 inversions and 1,051 translocations events for the rearrangement events that occurred between JD17 and WM82 (Table S7). Similarly, 86 and 1,273 inversions and translocations occurred between JD17 and ZH13, and 93 and 2,996 inversions and translocations occurred between JD17 and W05, respectively. We observed five large inversions (>1Mb) on Chr04, Chr05, Chr06, Chr07, Chr19 and two large translocation events (>200kb) on Chr02 and Chr07 (Figure S12-S14, Table S8), respectively, that were present in JD17 compared to all other three species. Examination of the breakpoint loci of these variants by comparing the sequenced subreads to the genome showed that the assembly of JD17 was correct (Figure S15-S21). Although the breakpoint at JD17 (Chr06:32,732,757) is an N sequence (i.e. gaps) and the subreads at Chr07:27,723,877 are poorly supported, the corresponding breakpoint positions in the WM82 genome are both gaps. This suggests that these structural differences such as inversions and translocations are more likely to be true genetic variation in JD17 relative to the other three strains.

### Identified PAVs by comparison with WM82, ZH13 and W05

Comparison of the genomes of JD17 and WM82 revealed 20,984 JD17- specific fragments (total length: 13.13 Mb) and 22,635 WM82-specific genomic fragments (total length: 30.81 Mb) (Table S9). Similarly, comparing the genomes of JD17 and ZH13 revealed 22,818 JD17-specific genomic fragments (13.68 Mb) and 23,456 ZH13-specific genomic fragments (33.27 Mb). Comparison of the JD17 and W05 genomes also revealed 36,658 JD17- specific genomic fragments (21.86 Mb) and 37,443 W05-specific genomic fragments (46.92 Mb). We further merged PAV sequences within 100 kb from the physical coordinates to identify PAV clusters (Table S10). The majority of these PAV sequences (99.8%) were shorter than 5 kb. We identified 23 PAV sequences longer than 5 kb in JD17 relative to Wm82, respectively. These PAV sequences were unevenly distributed in the genome (Figure 1), with some located in clusters. The largest of these PAV cluster fragments is a 1.6- Mb JD17-specific fragment containing 5 predicted genes located between 14.8 Mb and 16.4 Mb on chromosome 1. However, there is a PAV cluster of less than 0.7 Mb that is very rich in genes, up to 73 coding genes, located between 45.3 Mb and 46.0 Mb on chromosome 8. After GO enrichment analysis, they were found to be associated with NADH dehydrogenase, aspartate-type endopeptidase activity, histidine, protein and sugar de metabolism. (Figure S22)

Based on the criterion that a gene can be designated as a PAV gene if its coding sequence is covered by ≥ 75% of the PAV sequence (Sun et al. 2018), we compared JD17 with WM82, respectively, and we identified 100 JD17- specific and 75 WM82-specific PAV genes. Similarly, by comparing JD17 with ZH13, we identified a total of 80 JD17-specific and 156 ZH13-specific PAV genes, and between JD17 and W05, 101 JD17-specific and 67 W05-specific PAV genes were identified. It can be seen that there are large differences in genomic sequences between JD17 and WM82, ZH13 and W05.

By comparing these JD17-specific genes, it was found that only 20 JD17- specific genes were identified simultaneously (Figure S23), and most of the JD17-specific genes could still be found in different strains. The functional enrichment analysis of these 20 genes showed that these genes were related to signal transduction, response to stimulus, protein synthesis, and nitrogen metabolism. The results of functional annotation also showed that these genes were associated with powdery mildew and TMV resistance protein synthesis.(Figure S24)

### Identified SNPs and Indels by comparison with WM82, ZH13 and W05

To find SNPs and Indel between JD17 and the other three strains, we combined MUMmer and SyRI tools. The results showed that we identified 1,695,741 (2,675,463 in ZH13 and 3,664,629 in W05) SNP, 213,509 (324,109 in ZH13 and 391,977 in W05) insertions and 233,180 (391,977 in ZH13 and 408,512 in W05) deletions variations In JD17 genome by comparative genome-wide analysis. Combined with the results of structural variation and PAVs, both showed the smallest difference between JD17 and WM82 and the largest difference with wild-type soybean W05. This may be due to the fact that JD17 has the lineage of WM82, but it also supported that wild soybean can provide richer genetic diversity.

Annotation of these variants showed that they mainly affect intergenic regions, but there are still 4,947 variants between JD17 and WM82 that have a very large impact on 1,785 protein-coding genes. Enrichment analysis of these affected genes revealed that the primary function of these genes is associated with sulfate transmembrane transport, regulation of organ growth, regulation of developmental growth, carbohydrate binding. (Figure S25)

Similarly, a large effect of 7,363 variants on 2,567 protein-coding genes was observed between JD17 and ZH13, and a large effect of 6,674 variants on 3,611 protein-coding genes was observed between JD17 and W05.

### Identification of SNF genes and differences among JD17, WM82, ZH13 and W05

To identify genes associated with SNF in the four genomes, we blast identified genes from published articles to these genomes. We identified 331, 360 and 285 and 346 genes potentially related to nitrogen fixation in JD17, WM82, ZH13 and W05 genomes, respectively (Table S11). Of these, 88.8% of the SNF genes were identified in all four soybeans, and in JD17, 96% of the genes were identified, while fewer genes were identified in ZH13 (Figure S26). Subsequently, by comparative analysis of SNF genes from these four species, there are two expansion genes (*LjCLE-RS2* and *MtNAC969*), two contraction genes (*LjCLC1* and *MtCP6*), and five lost genes (*GmENOD93*,*GmRj2_GmRFG1*,*LjENOD40-1/ENOD40-*2,*LjHIP*,*MtCAS31*) (see Table 12 for details) in JD17 relative to their common ancestor (Table S12). By examining the structural variation of 331 genes potentially associated with nitrogen fixation in JD17, we finally found 18 genes with structural variation between WM82, ZH13 or W05 and affecting protein coding (Table S13). Further by constructing an evolutionary tree to manually confirm the effect of these genes undergoing structural variation on them, we finally observed that nine of them were not in the same branch of the evolutionary tree as the published sequence of WM82. For example, the *LjIGN1* genes (Glyma.10G291900 and Glyma.20G241200), which encodes the synthetic Ankyrin-Repeat Membrane Protein, which is important for symbiosis and nodulation(Kumagai et al. 2007). By evolutionary analysis of their homologous genes, we found that Glyma.10G291900 and Glyma.20G241200 are located in two branches, and the branch where Glyma.10G291900 is located has corresponding genes in all four species with essentially the same protein code (Figure S27). However, the branch of Glyma.20G241200 showed differences. By multiple sequence comparison, JD17020G0232500 in fact encodes 192 amino acids that are very different from several other proteins (Figure S28), and this genetic difference most likely affects symbiosis and nitrogen fixation and still need further studies to be verified.

## Conclusions

Here, we constructed a high-quality de novo assembly of the soybean cultivar Jidou 17 (JD17) with high continuity, completeness, and accuracy, with contigs N50 is close to 18 Mb. We predicted 52,840 gene models and reconstructed 74,054 high quality full-length transcripts. By performing a genome-wide comparative analysis of the reference genome of JD17 with three published soybean (WM82, ZH13 and W05), we identified five large inversions and two large translocations, and 20 PAV genes, which are specific to JD17. SNPs and Indels between JD17 and the three soybeans were also identified and annotated separately. In total, 1,785 genes in JD17 were found to have altered protein coding due to these structural variants, which may affect their function. Further identification of SNF genes and assessment of the impact by these variants revealed that the coding sequences of 9 SNF genes were significantly affected. We hope that our dataset will provide a valuable resource for comparative genomics and functional genomics of soybean.

## Data availability

The sequencing data (PacBio Whole Genome Sequencing data for assembly, resequencing data, Hi-C data and Mixed-Sample RNA-seq data for Annotation) used in this study have been deposited into National Center for Biotechnology Information under BioProject Number PRJNA412346 with accession number SRR12416523 - SRR12416632, SRR12416444, SRR12416750 and SRR9643849 - SRR9643851. The final assembly had been deposited at GenBank JACKXA000000000.

## Author information

### Contributions

NC, JZ and XW coordinated the research. XY, SC, HW, LL, BA and WF prepared the DNA and RNA samples for sequencing. SL, DK and RW performed PacBio sequencing. XY, JL, HW, ML, QU and LL performed assembly analysis and annotation of the genome and transcriptome. CY and MZ provided the plant seed from Hebei Academy of Agricultural and Forestry Sciences. XY, JZ, NC and XW wrote the manuscript. All authors have read and approved the final manuscript.

---

**Table S11.**
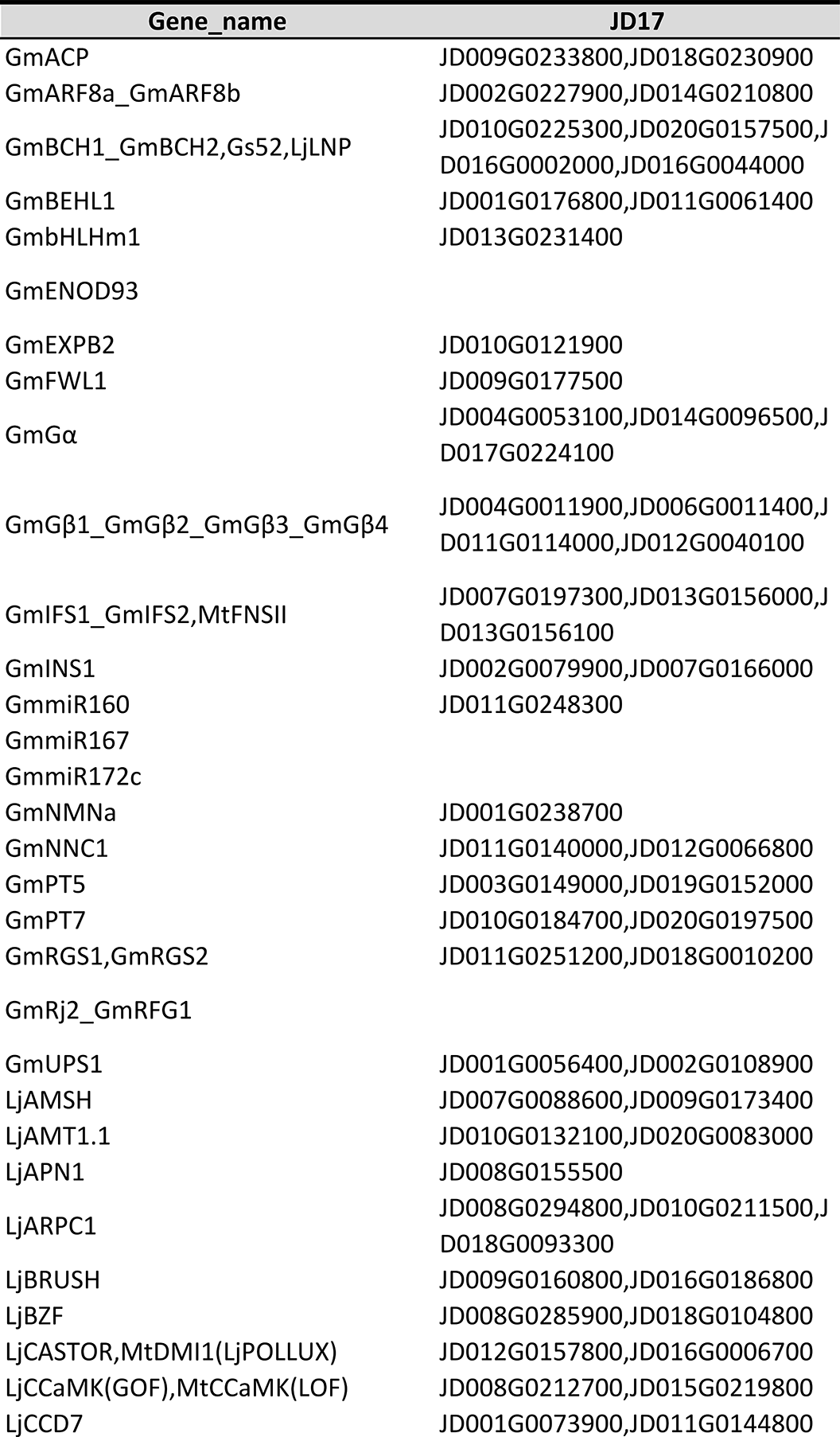
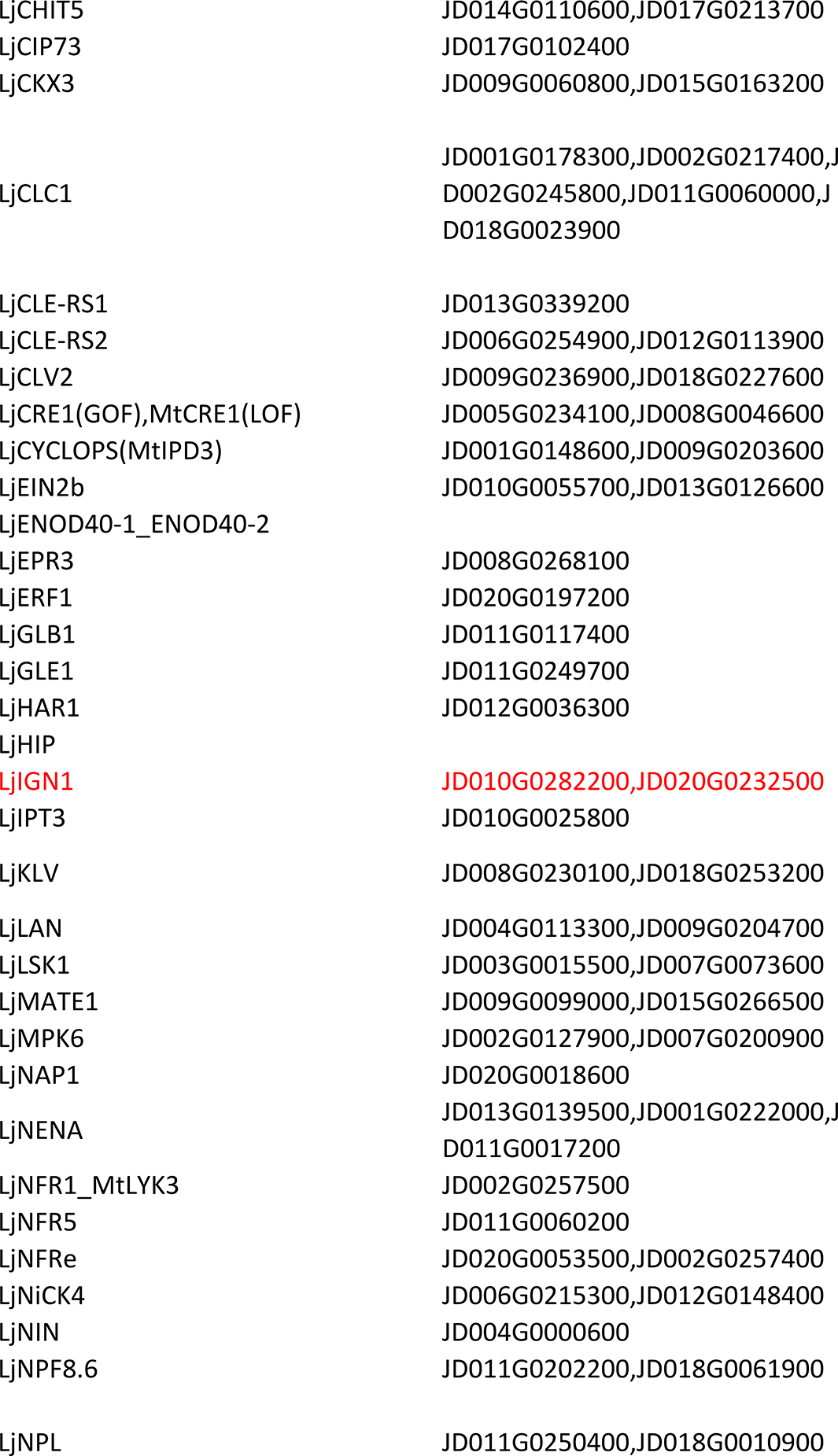
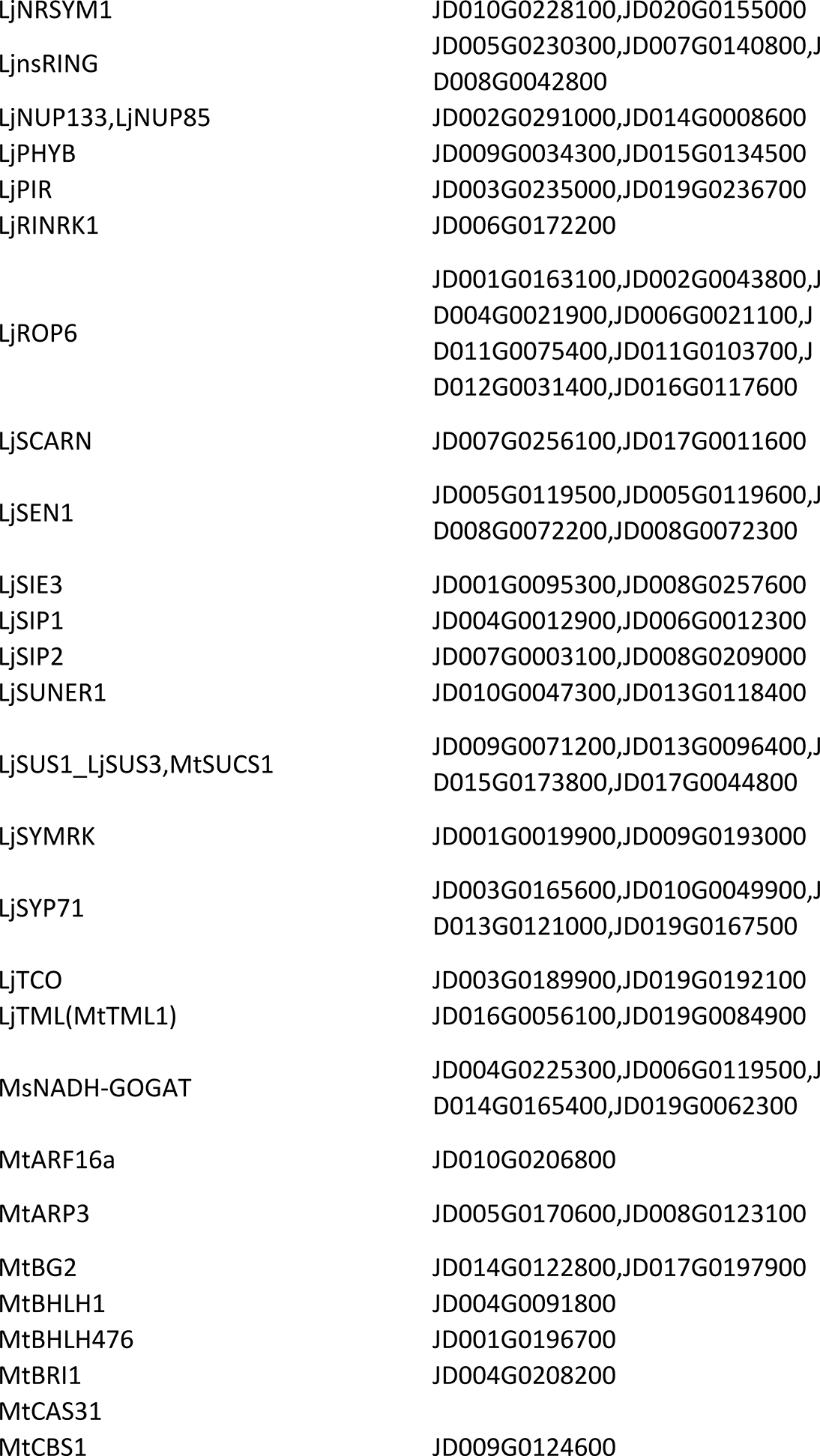
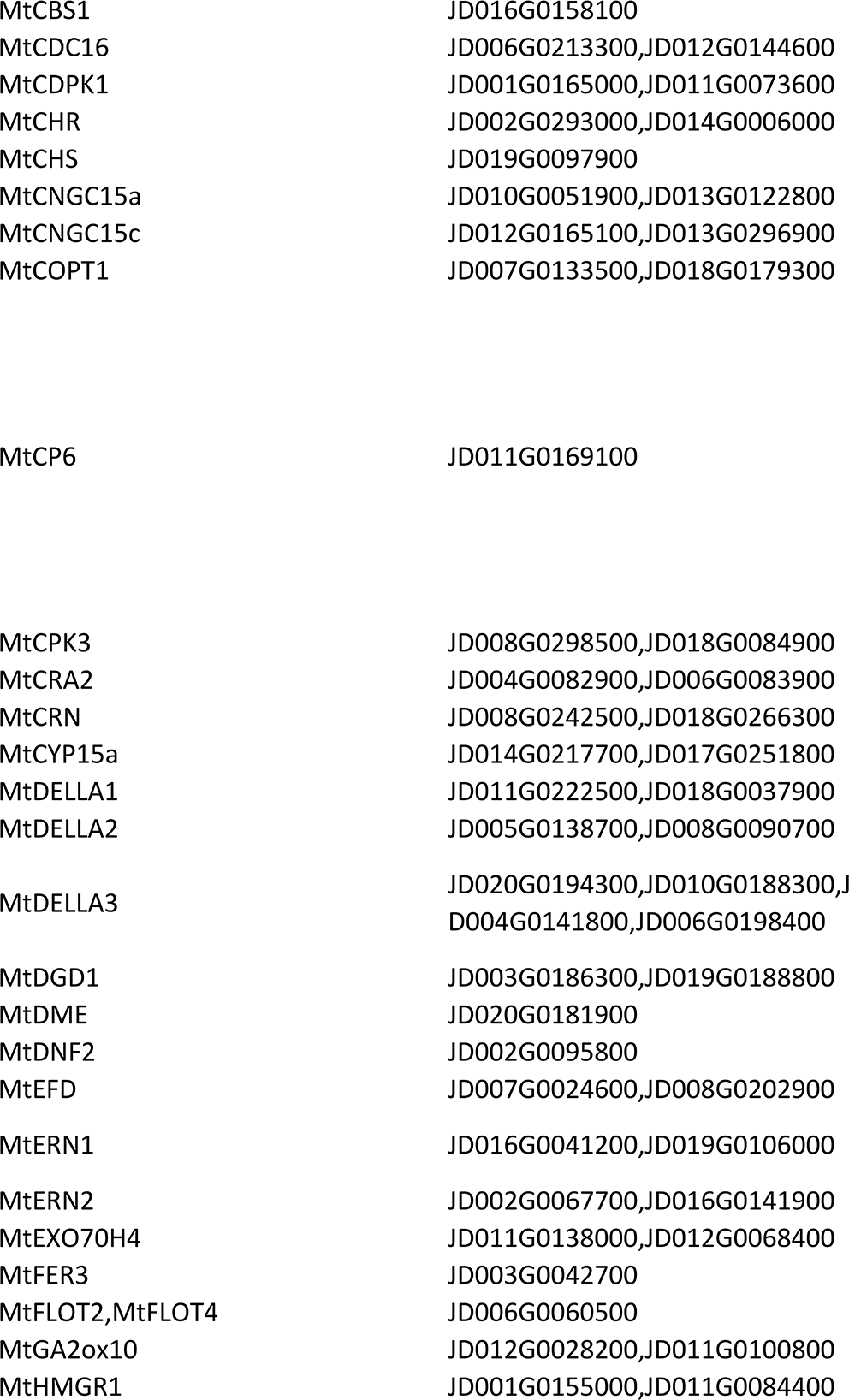
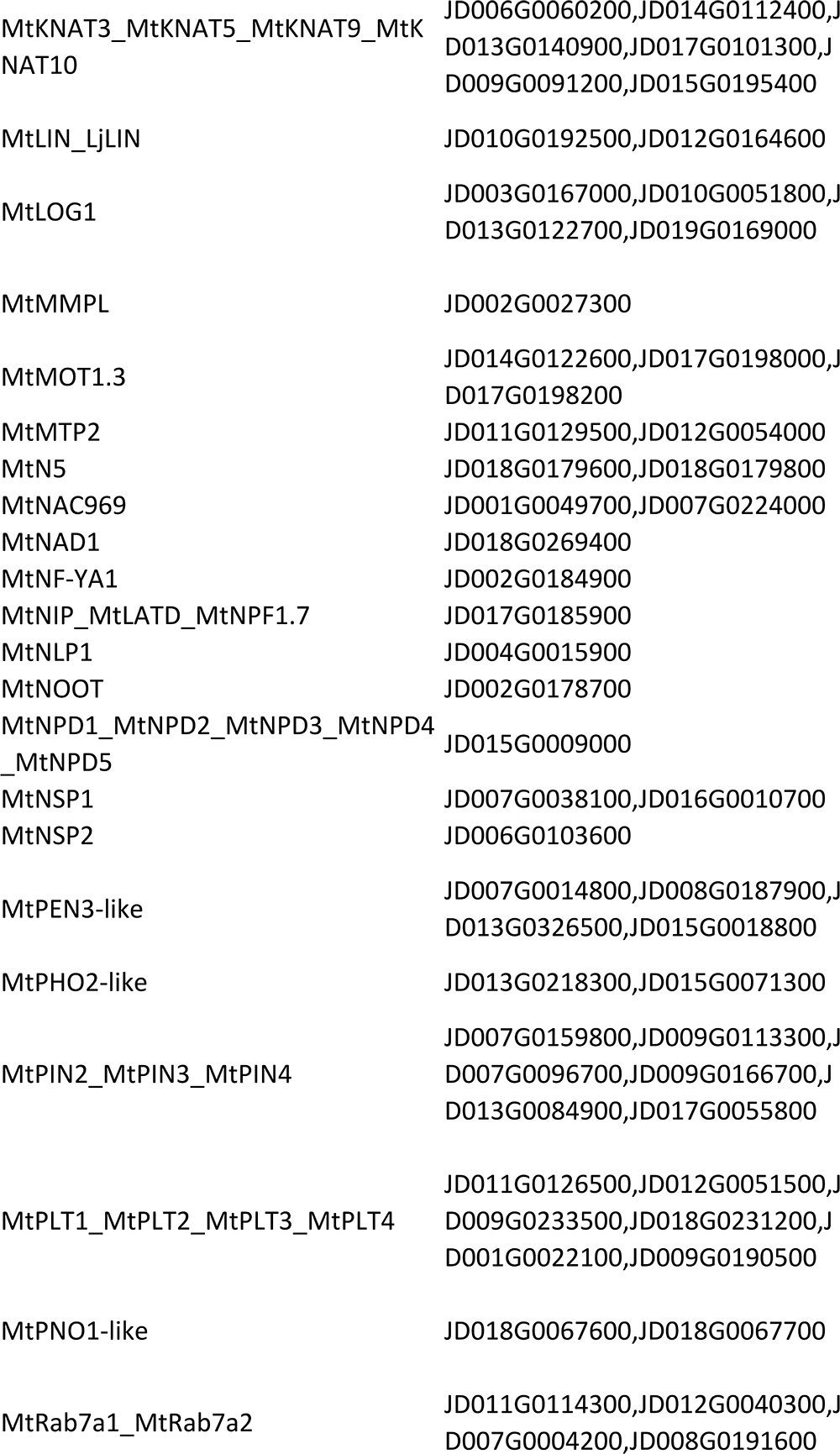
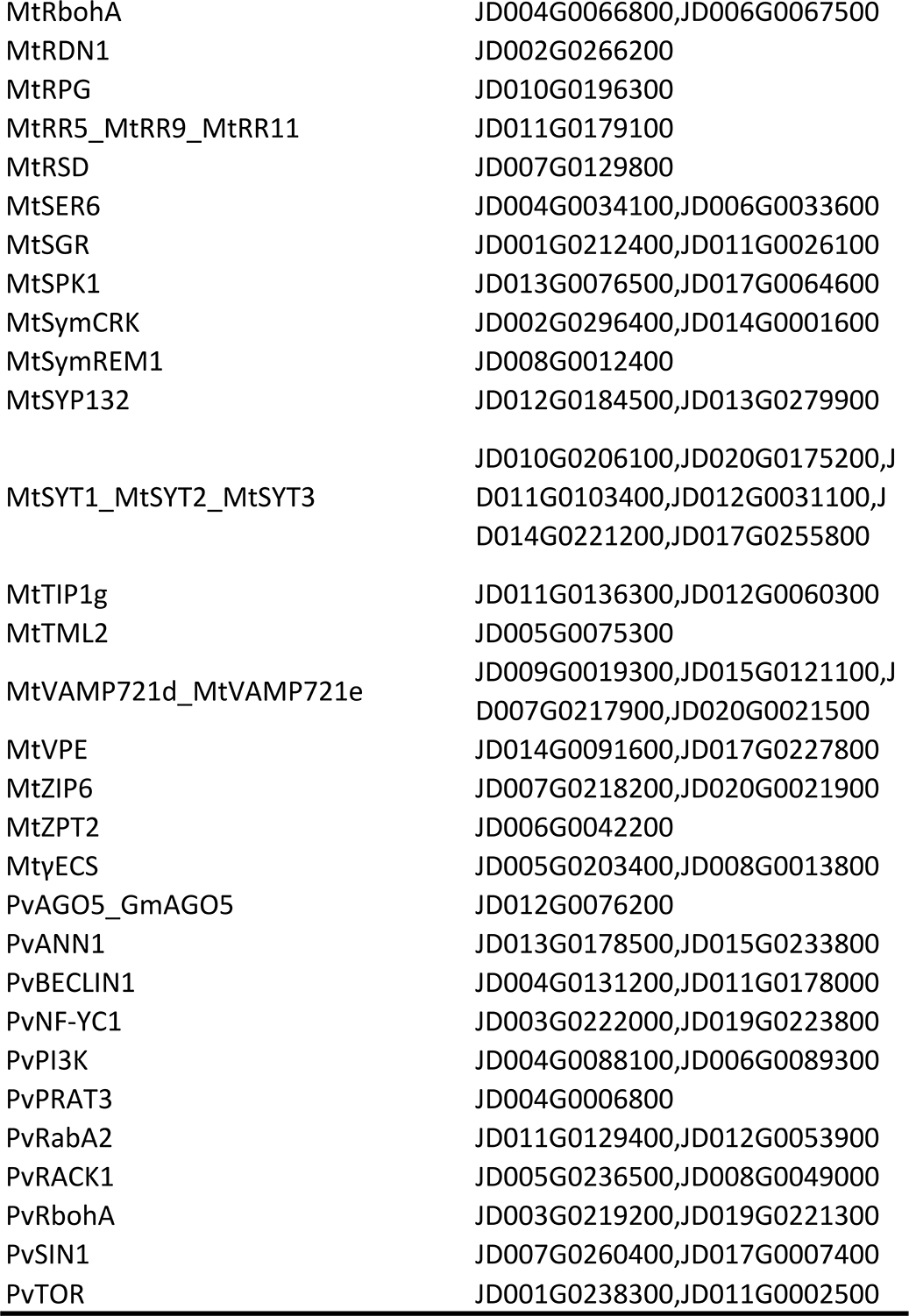
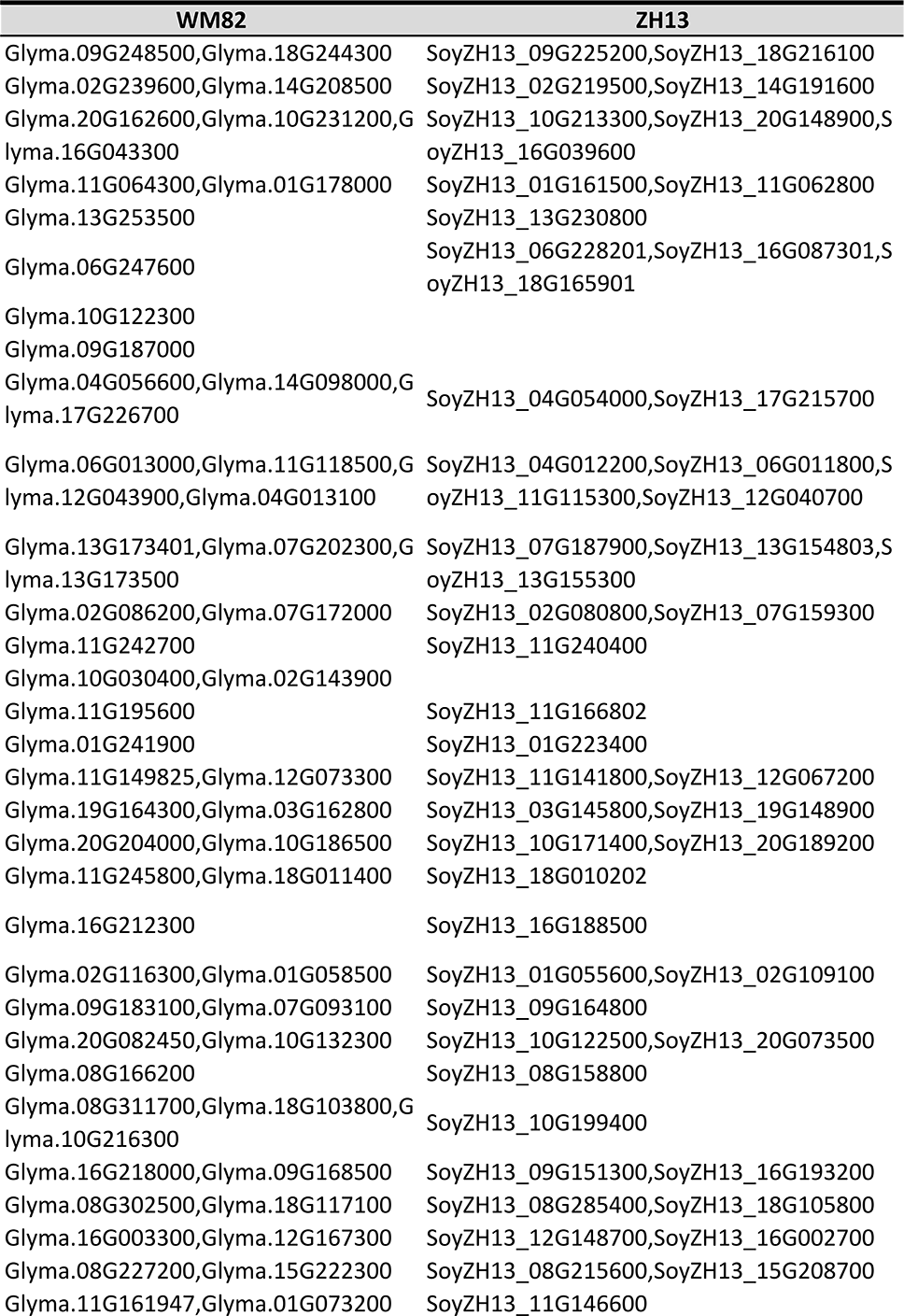
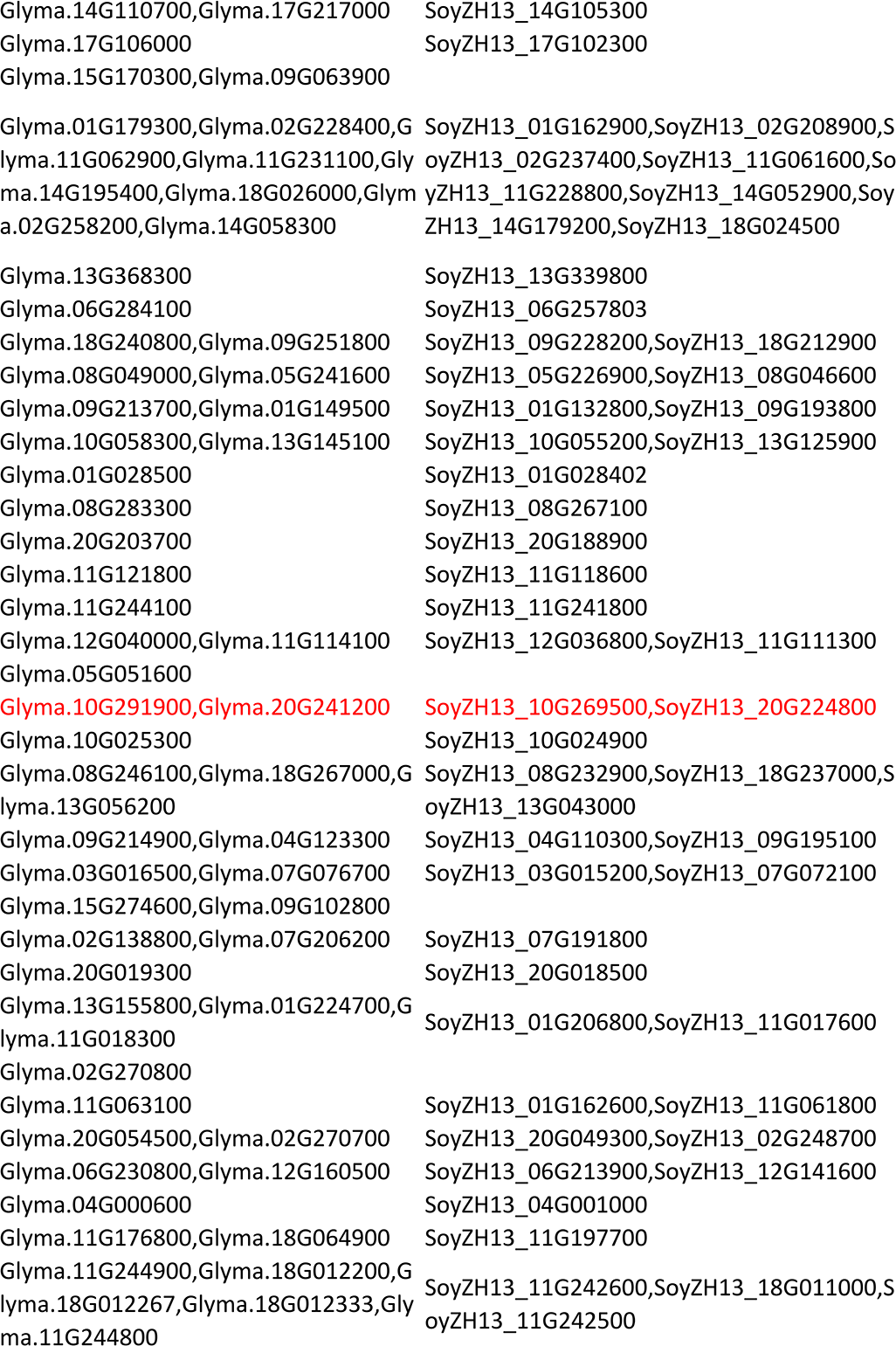
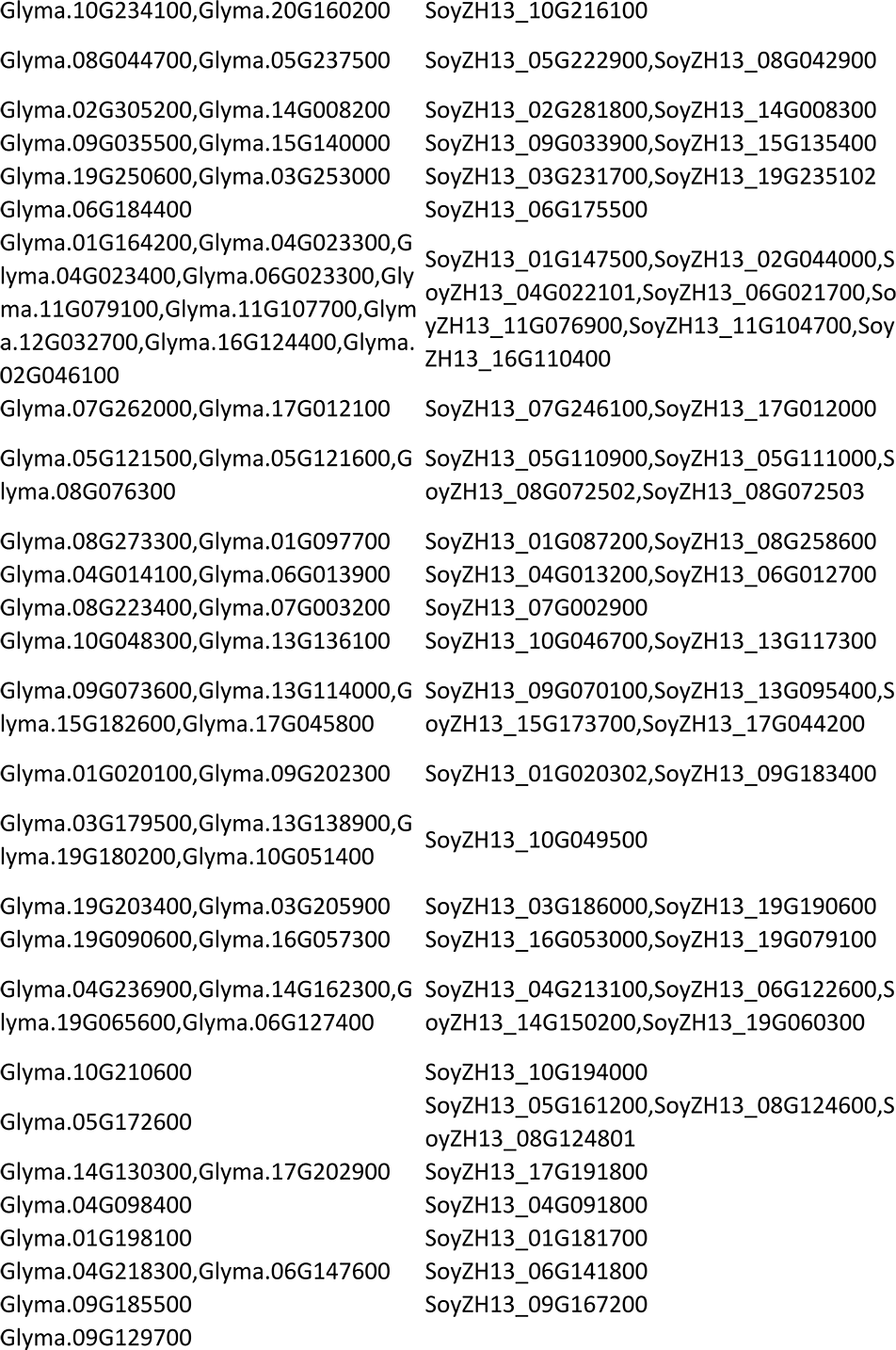
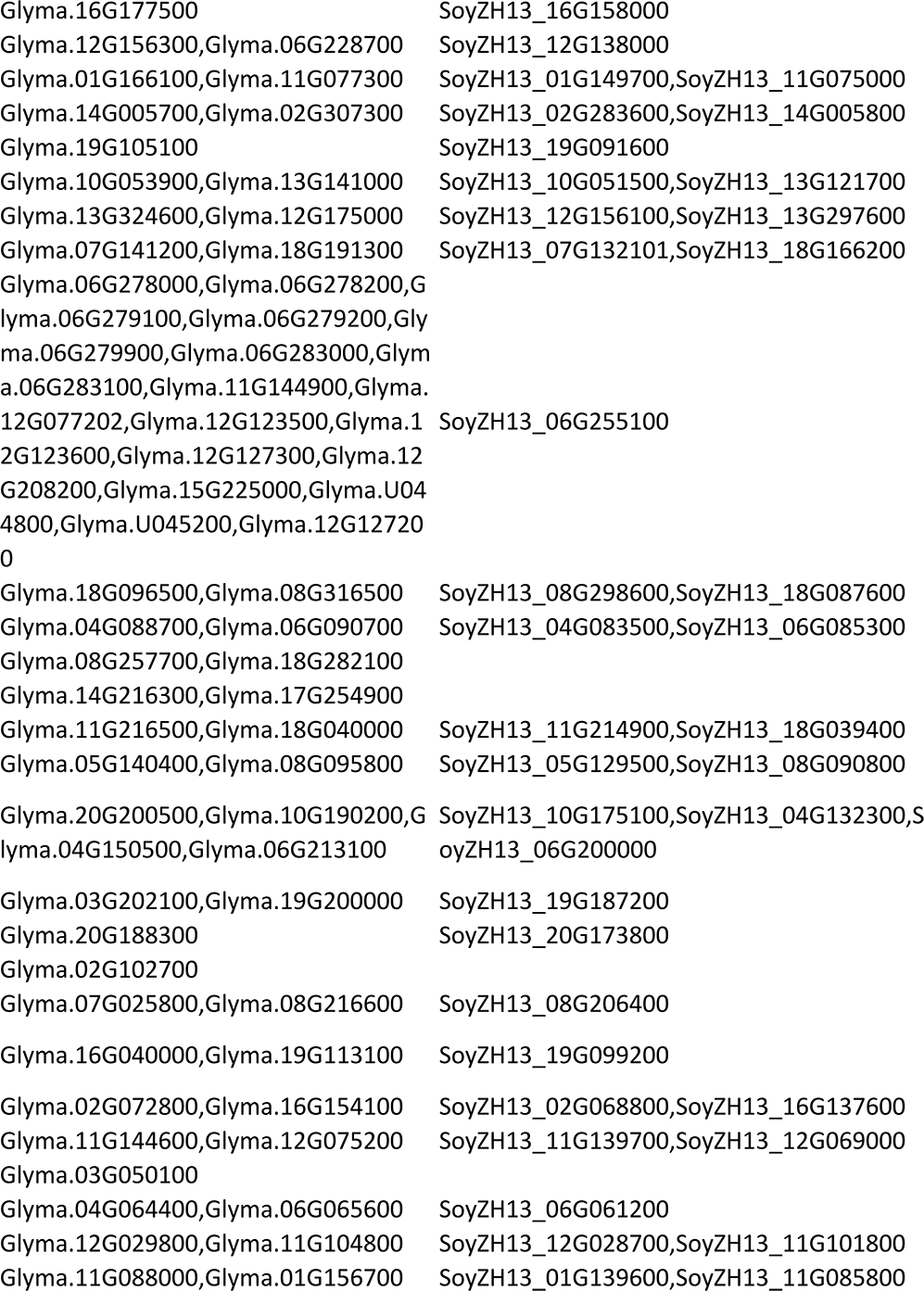
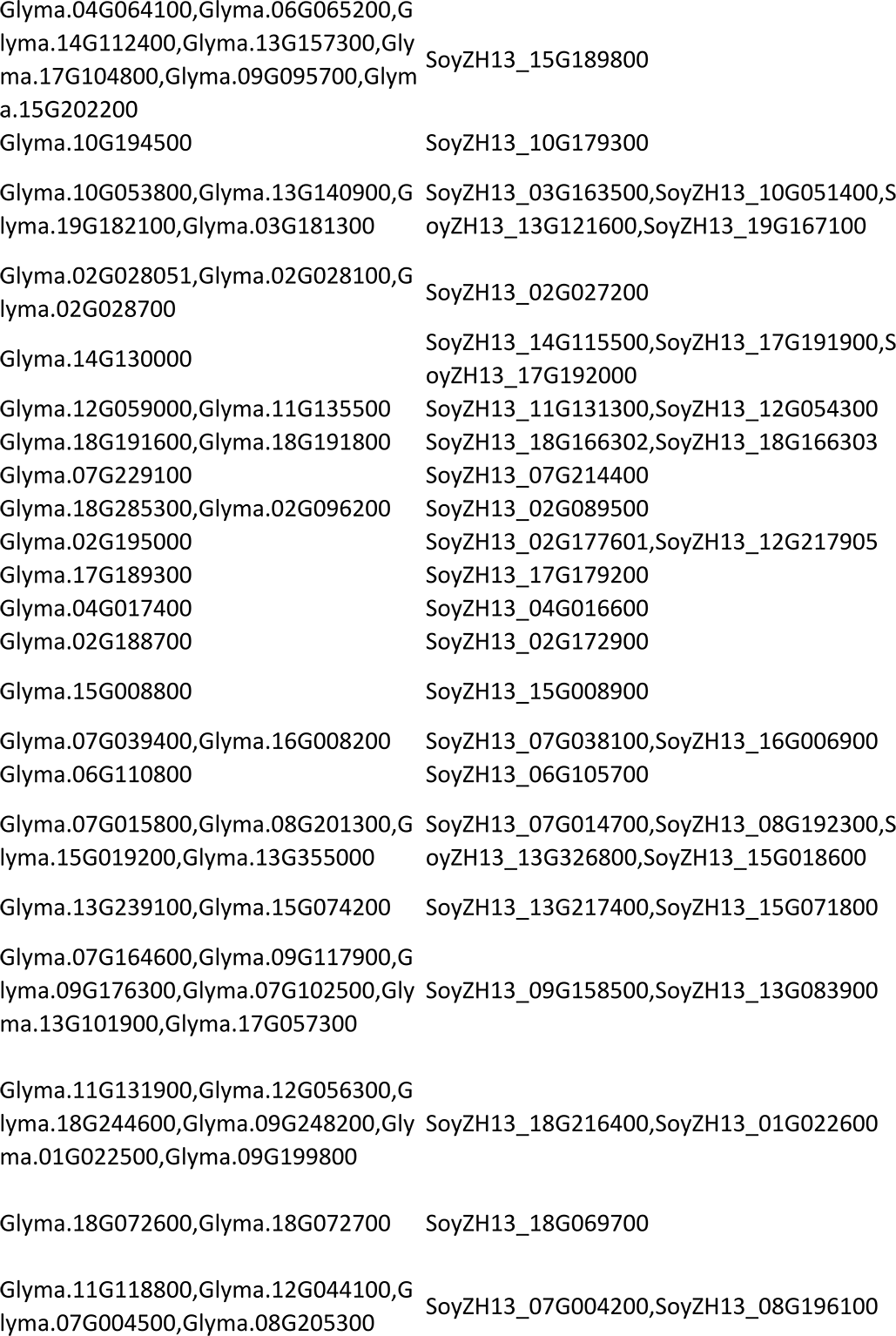
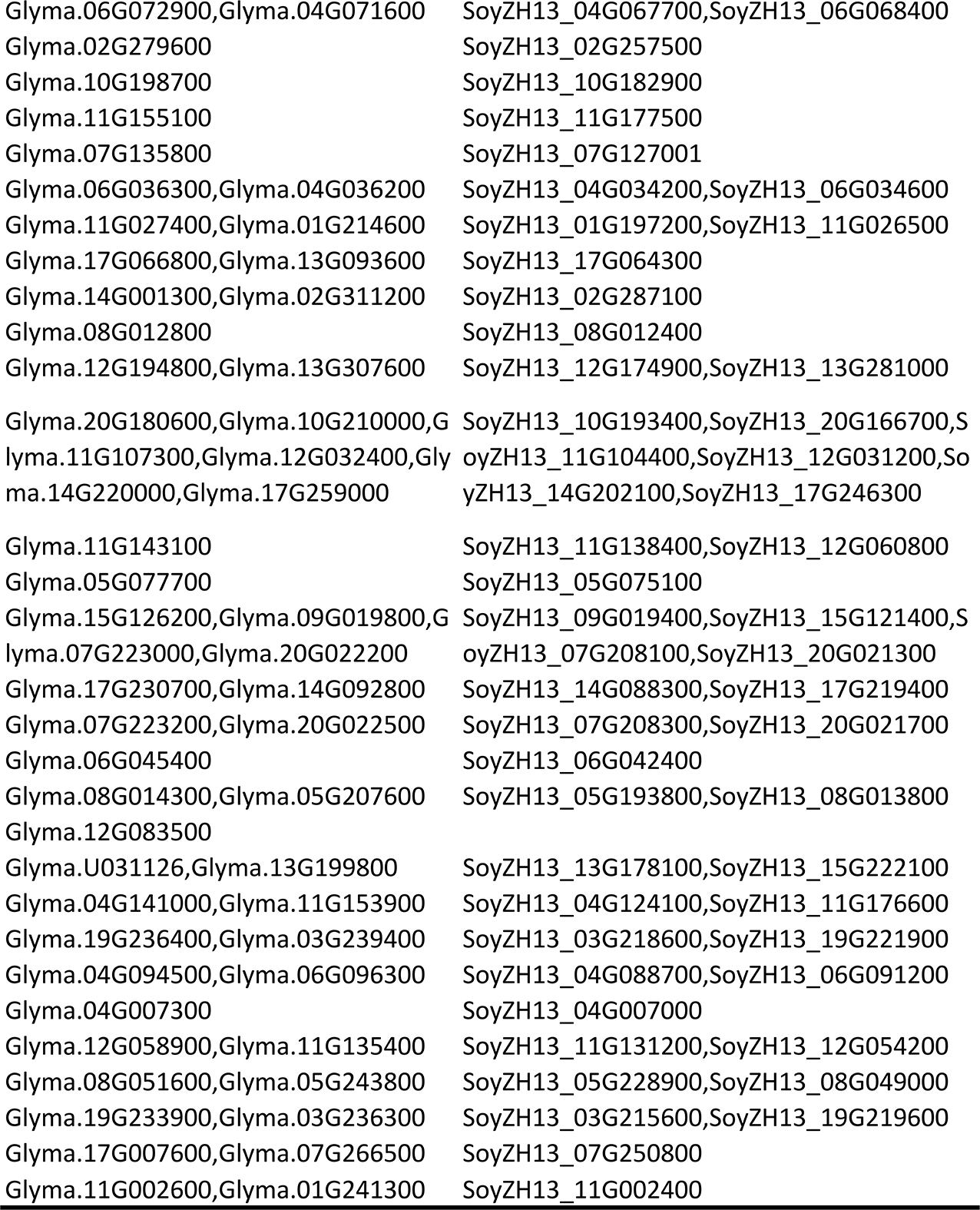
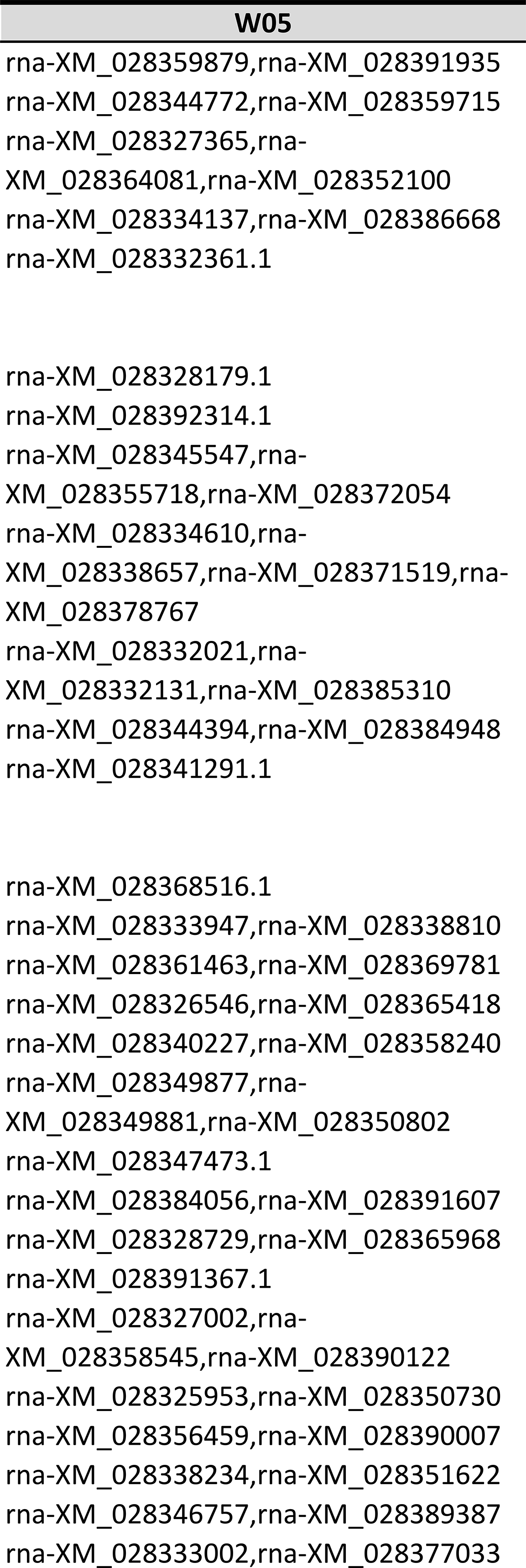
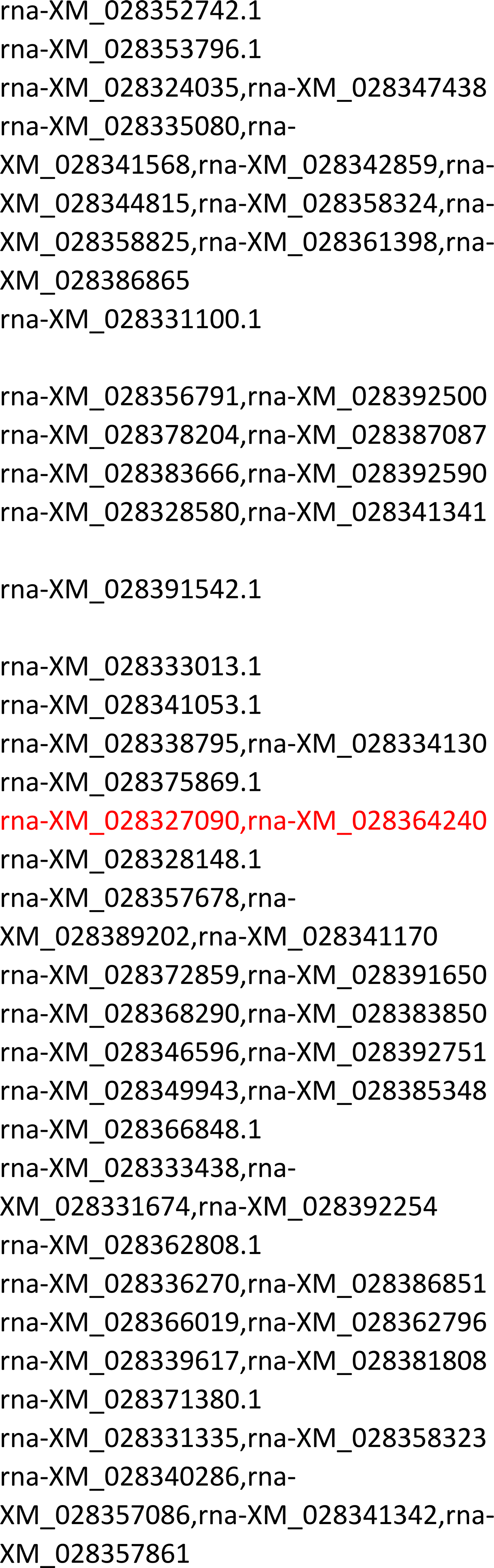
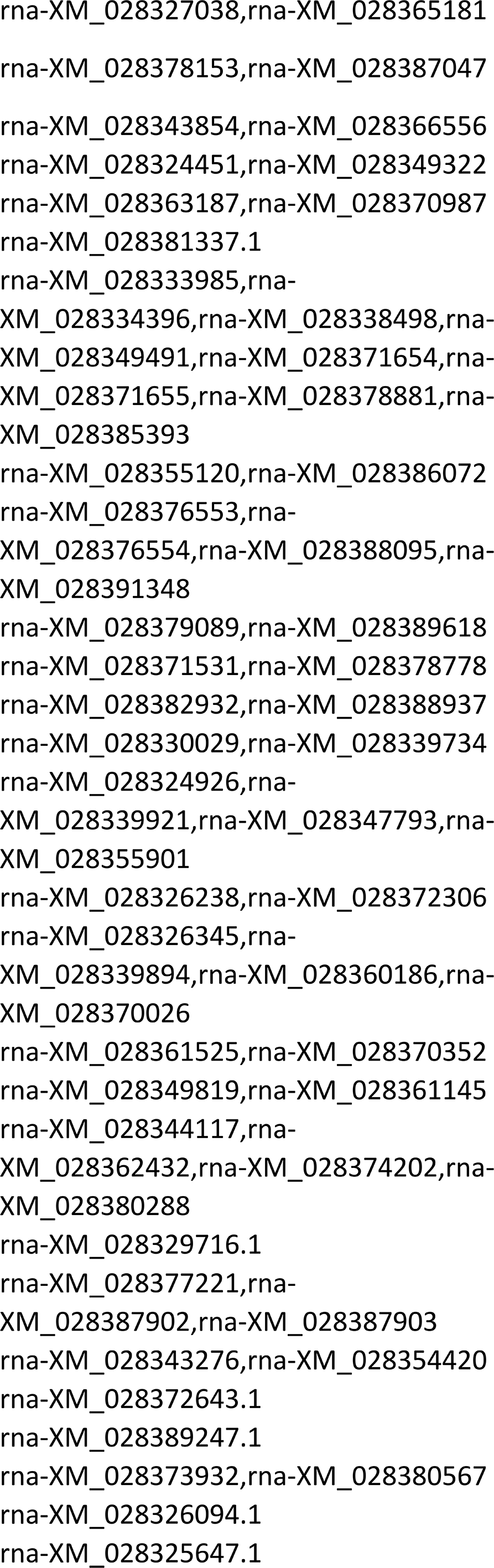
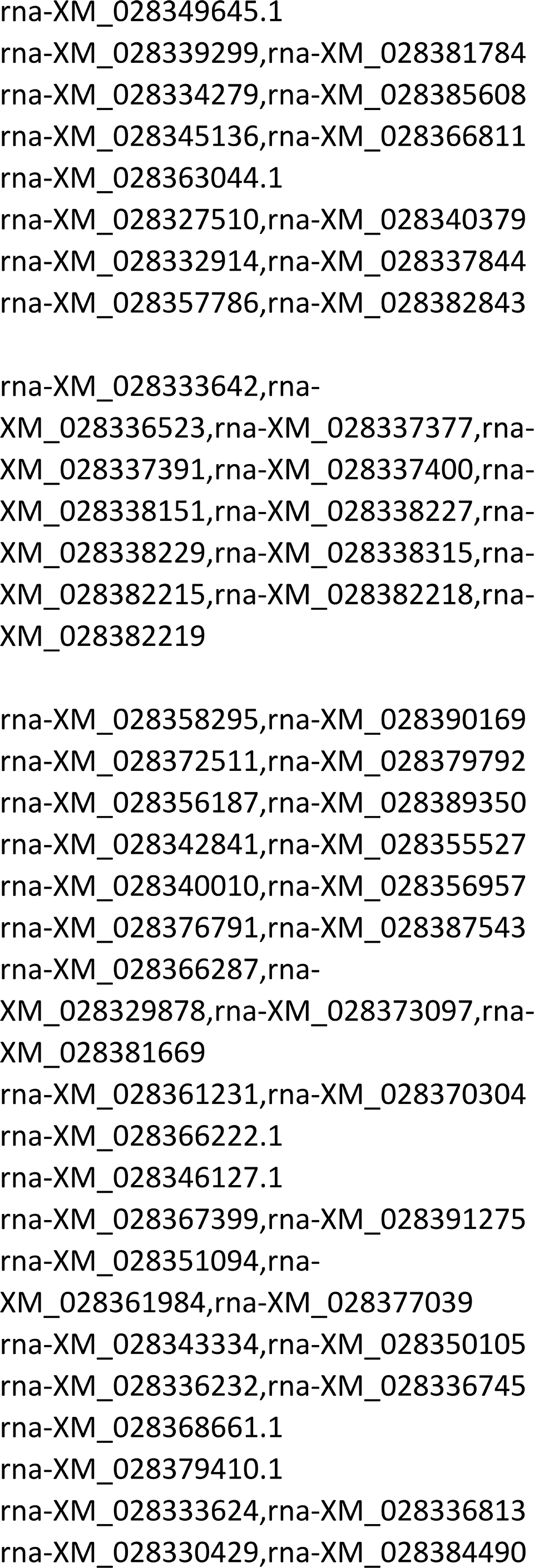
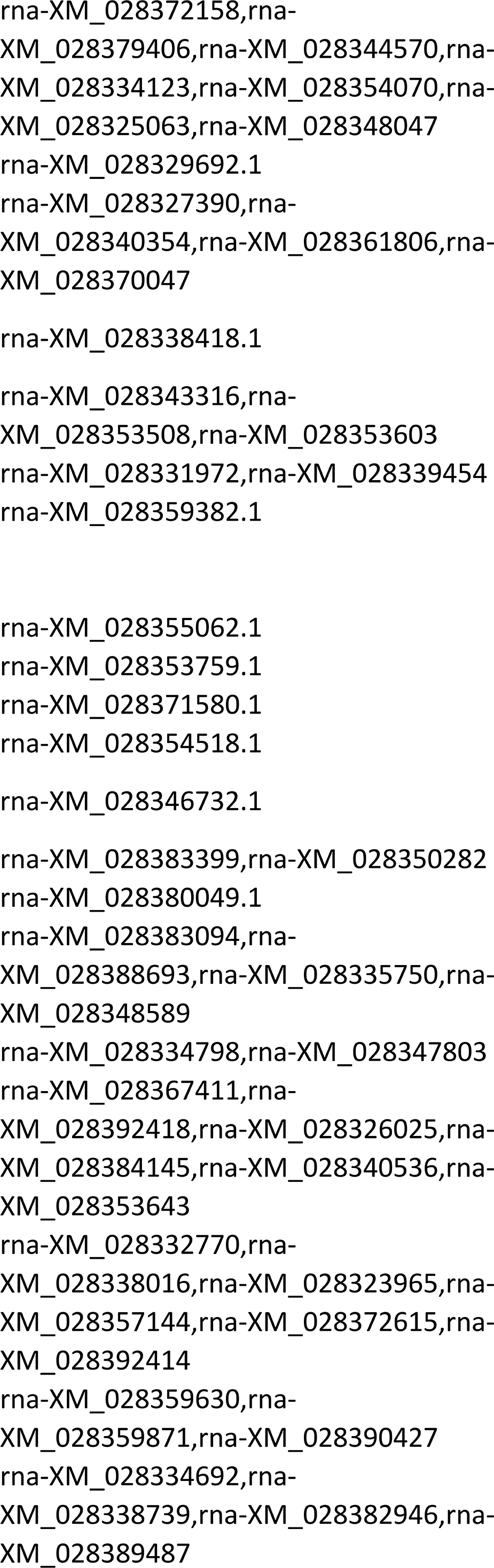
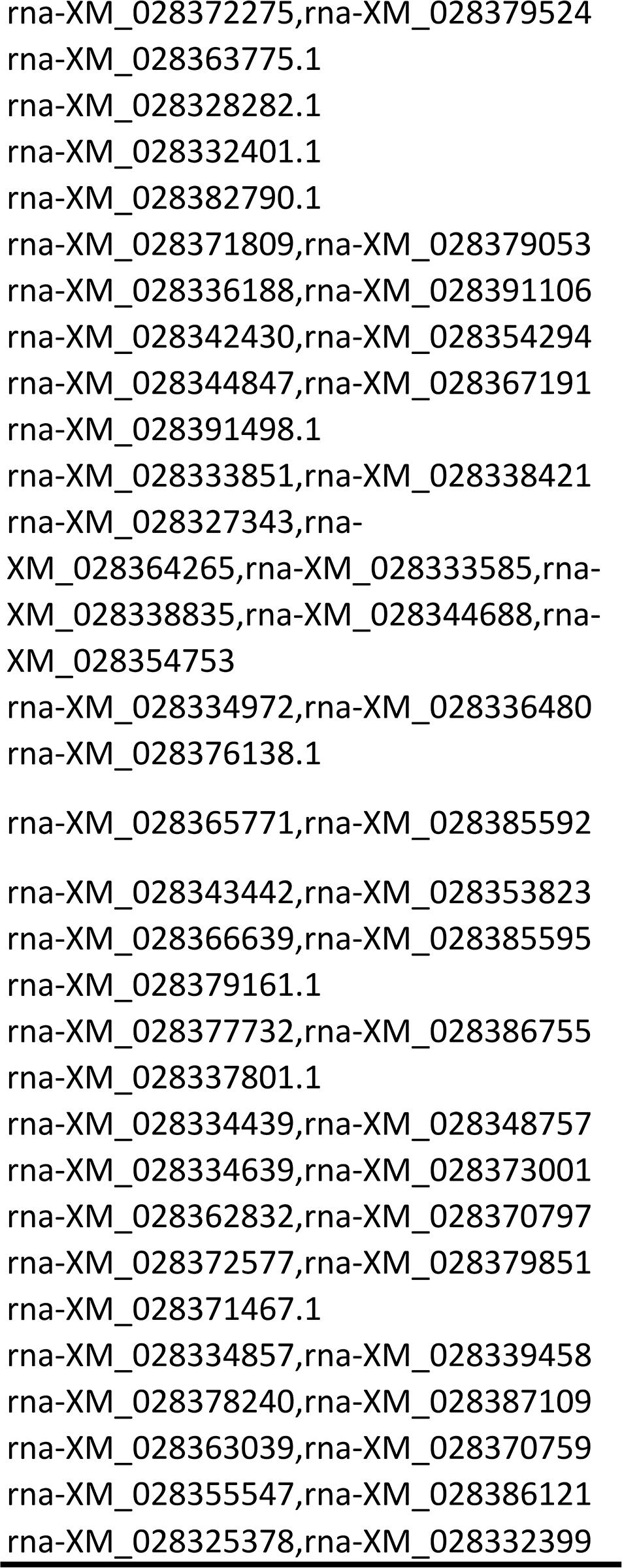
SNF genes identified in JD17, WM82, ZH13 and W05.

